# Long-Term Rock Dove (*Columba livia*) Primordial Germ Cell Culture: A Tool for Avian Conservation

**DOI:** 10.1101/2025.09.17.676777

**Authors:** Martin W. Nicholson, Lucas Moreira, Emily A. Boyle, Evan McCabe, Adrianna Soriano, Malcolm McSwain, William N. Feist, Samantha K. Nguyen, Kaitlin Steiger, Nicolas Alexandre, Savannah J. Hoyt, Nicole M. Tillquist, Rachel J. O’Neill, Kennosuke Ichikawa, Mike J. McGrew, Olivier Fedrigo, Anna Keyte

## Abstract

Primordial germ cells (PGCs) are critical tools for genome engineering and conservation in birds. While culture systems for chicken PGCs have been well established for nearly two decades, efforts to propagate PGCs from other avian species have proved exceptionally challenging, limiting the broader application of artificial reproductive technologies in birds. Here we report the first successful derivation and long-term culture of PGCs from the rock dove, or common pigeon (*Columba livia*). Guided by transcriptomic profiling of PGCs, we developed a species-specific medium that supports PGC maintenance and expansion. We identify insulin signaling as essential for survival and demonstrate that retinoic acid receptor inhibition, while maintaining vitamin A, is necessary for propagation. Supplementation with BMP4, LIF, GDNF, and pleiotrophin further enhances PGC proliferation. Cultured cells express canonical germline markers and migrate to the gonads following injection into both rock dove and chicken embryos, confirming functional competency. These findings establish a platform for germline manipulation and biobanking in Columbidae, broadening the applicability of reproductive technologies to conservation efforts.

## Introduction

Primordial germ cells (PGCs) are the embryonic precursors to gametes—eggs and sperm—and are essential for the transmission of genetic material across generations. In birds, PGCs arise early in development in the central region of the blastodisc, then translocate to the anterior germinal crescent and enter the embryonic bloodstream, migrating to the gonadal ridges where they colonize the developing gonads and differentiate into spermatogonia or oogonia (Swift 1914; Ginsburg 1994; De Melo Bernardo et al. 2012). This distinctive migratory behavior allows PGCs to be isolated from multiple developmental stages, including the embryonic bloodstream and gonads (Chen et al. 2020; Clawson and Domm. 1969; Whyte et al. 2015; van de Lavoir et al. 2012). Combined with their early segregation from somatic lineages, this accessibility makes avian PGCs an invaluable system for studying germline development and reproductive biology.

In addition to their utility as a model, avian PGCs have become central to applied germline manipulation strategies. In contrast to mammals, where cloning and somatic cell nuclear transfer enable genetic modification, such methods are not tractable in birds. As a result, PGC-mediated germline transmission has emerged as the primary platform for avian genetic engineering and biobanking—two critical tools for conservation and synthetic biology (Han and Park, 2018; Ichikawa and McGrew, 2024).

Significant progress has been made over the past two decades in developing protocols for the *in vitro* maintenance, genetic modification, and germline transmission of chicken PGCs (van de Lavoir et al. 2006; Whyte et al. 2015). Cultured chicken PGCs can be genetically engineered and reintroduced into host embryos to generate germline chimeras, facilitating applications in developmental biology, agriculture, and biotechnology (Taemeh et al. 2025; Tajima et al. 1993; Woodcock et al. 2019). Importantly, this system also enables cryopreservation and long-term storage of germline cells for biobanking purposes—an urgent priority in the face of accelerating biodiversity loss (Hu et al. 2022; van Oosterhout et al. 2025; van de Lavoir et al. 2012). However, the application of PGC-based technologies has remained largely restricted to domestic chickens (*Gallus gallus*), and efforts to translate these approaches to other bird species have been met with limited success.

PGC culture systems have recently been adapted for other species. For example, feeder-free media have been developed for chicken PGCs using combinations of fibroblast growth factor (FGF), activin A, and bone morphogenetic protein 4 (BMP4) signaling, while goose PGCs have been shown to depend on cholesterol, FGF, and BMP4 signaling (Doddamani et al. 2025). Notably, activin A inhibits proliferation in goose PGCs—highlighting the species-specific nature of PGC signaling requirements. While there are reports of PGC culture for quail, duck, and zebra finch, these systems support only short-term proliferation (up to ∼50 days) (Kim et al. 2005; Park et al. 2007; Yakhkeshi et al. 2018; Gessara et al. 2021).

In the case of the rock dove, or common pigeon (*Columba livia*)—one of the most widely distributed birds globally—no protocols have been established for the derivation or extended culture of PGCs. This gap presents a major limitation for the application of artificial reproductive technologies and germline biobanking within the Columbidae family, which includes many species of conservation concern (Walker 2007).

In this study, we address the critical need for species-specific approaches to avian PGC culture by establishing and validating a robust system for the derivation and long-term expansion of pigeon PGCs. Applying single-cell RNA-sequencing (scRNA-seq), we characterize the molecular and cellular requirements necessary for long-term pigeon PGC culture. We identify previously uncharacterized dependencies on insulin signaling and retinoic acid receptor inhibition and define the contributions of specific growth factors to cell proliferation and maintenance of germline identity. Importantly, we demonstrate that cultured rock dove PGCs retain the capacity to migrate to and colonize the gonads of host embryos. These findings represent the first successful derivation and long-term culture of PGCs from a Columbidae species and provide a framework for the extension of PGC-based technologies to a broader array of avian species. Our findings not only expand the toolkit for studying avian developmental and synthetic biology but also provide a foundational method for avian germplasm preservation, with broad implications to the conservation of biodiversity of Columbidae.

## Results

### scRNA-Seq Informs Culture Optimization

We collected gonadal samples from six stage HH28 rock dove embryos, which are expected to contain undifferentiated PGCs (Hamburger and Hamilton, 1951). Samples were processed for scRNA-seq using Particle-templated Instant Partition sequencing (PIP-seq; Clark et al. 2023). Reads were mapped against the NCBI rock dove reference genome assembly and processed through the PIPseeker (Fluent Biosciences) pipeline. After stringent quality control (Supp. Fig. 1), our scRNA-seq dataset contained a total of 4,260 cells (Supp. Fig 2). We performed normalization and variance stabilization using the SCtransform method (Hafemeister and Satija, 2019), while regressing out known technical and biological confounders including cell cycle phase, mitochondrial gene content, and transcript count (nUMI). To characterize and quantify distinct cell populations, we applied unsupervised clustering on the dataset using UMAP for visualization, based on the expression profiles of 14,080 genes. This analysis revealed nine transcriptionally distinct clusters (Figure 1A). Cell type identities were assigned to each cluster based on the expression of canonical marker genes in the top 50 differentially expressed genes per cluster (Supp. Fig. 3). Gonadal PGCs (gPGCs) were easily identified as a single cluster of cells expressing canonical germline markers (e.g., *DDX4, DAZL, TDRD9, POU5F3/OCT4*) (Figure 1B).

**Figure 1:**
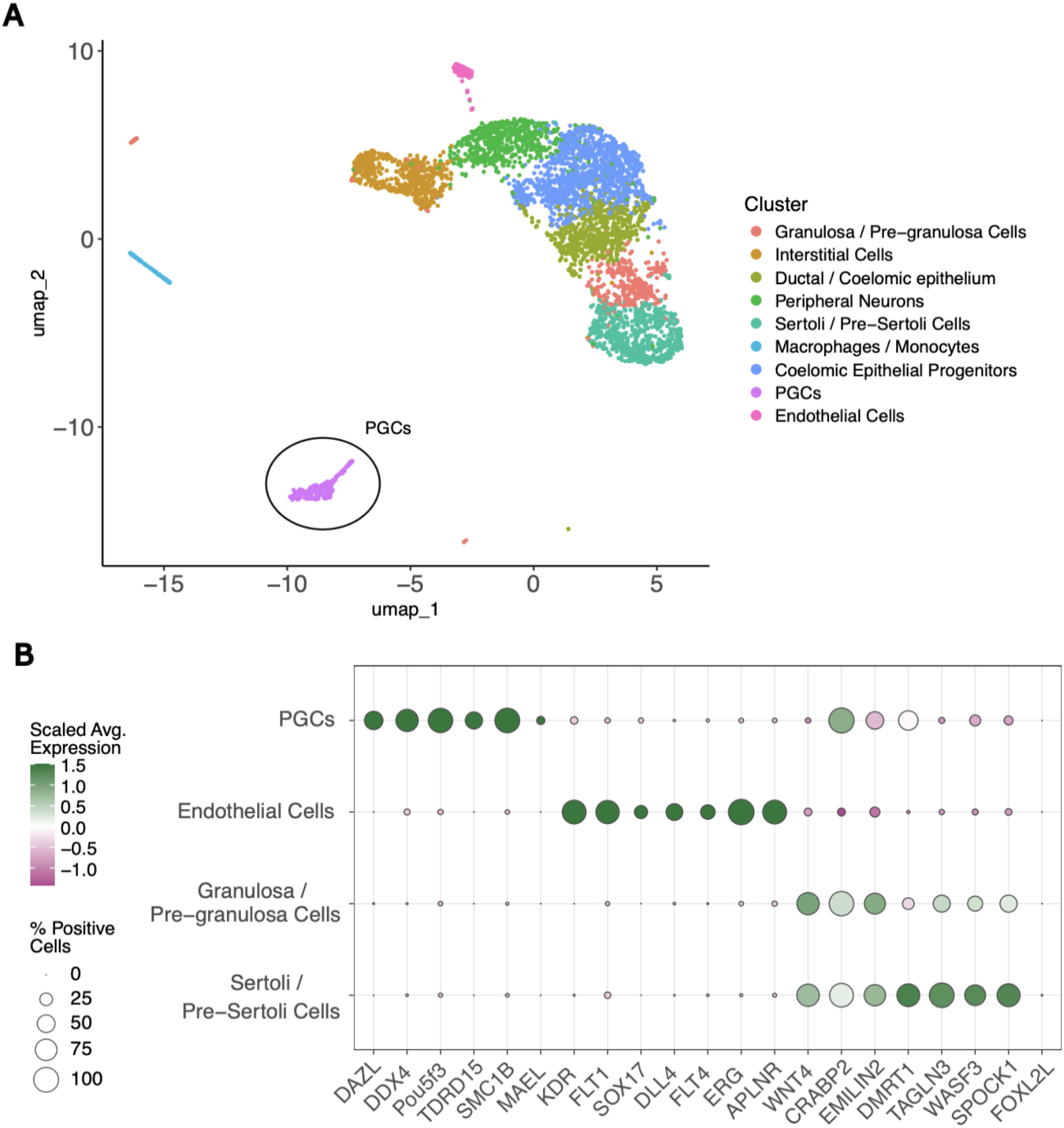
Clustering of rock dove scRNA-seq. ScRNA-seq clustering of rock dove gonadal cells. (A) UMAP of all cells identified in the gonadal tissue. (B) Expression patterns for canonical genes of the main cell types in rock dove gonads.

We then evaluated gene expression of cell surface receptors including those for fibroblast growth factor (FGF), insulin signaling, transforming growth factor (TGFβ), and bone morphogenetic protein (BMP) molecules, each of which has been shown to be involved in growth and survival of PGCs (Lochab and Extavour, 2017), as well as retinoic acid signaling pathway members, which have been implicated in germ cell differentiation (Anderson et al. 2008). We estimated receptor competence as the median fraction of PGCs expressing the receptors for a given growth/maintenance-related pathway. By this metric, IGF-family receptors show 49% competence (notably IGFR1) and FGF receptors show 50%. These levels are consistent with survival signaling via PI3K–AKT downstream of IGF and mitogenic signaling downstream of FGF. We also observed substantial competence for KIT/c-KIT (44%), the GDNF co-receptor complex GFRA1/RET (38%), and WNT pathway receptors (35%), implicating these pathways in PGC maintenance

and proliferation. By contrast, NOTCH and Hedgehog receptors showed comparatively low, patchy expression (19% and 21%), suggesting these common signaling routes are not primary growth drivers in rock dove PGCs (Figure 2A).

**Figure 2:**
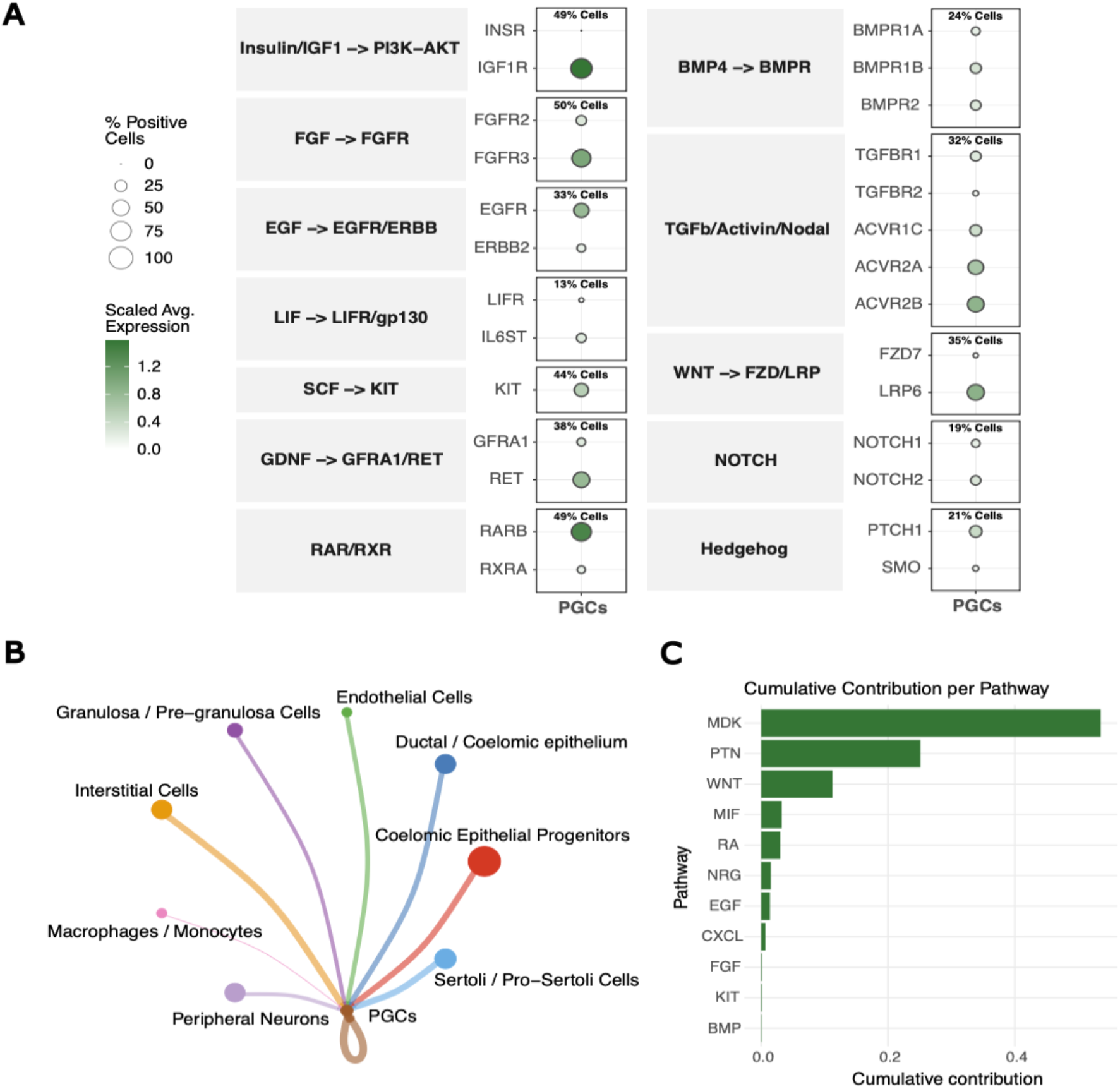
Pathway analysis of rock dove PGCs. (A) PGC cell surface receptor and pathway expression analysis. Receptor competence was calculated as the median percentage of PGCs expressing the receptors for a given growth/maintenance-related pathway. (B) Signaling pathway interactions identified with CellChat. Thickness of lines represent the strength (or relative probability/weight) of inferred communication between two cell types. (C) Cumulative contribution of growth-related pathways via PGC receptors.

While vitamin A (retinol) is required for membrane and metabolic homeostasis, its metabolite retinoic acid (RA) activates RAR/RXR to promote germ-cell differentiation and meiotic entry (Zhang et al. 2015; Tedsco et al. 2013). PGCs showed high RAR/RXR receptor competence (49%; notably RARB), implying that unopposed RA signaling could erode PGC proliferation and identity in culture (Figure 2A).

Since cellular growth and maintenance also depend on intercellular signaling, we also investigated gPGCs’ cell-cell communication. To this end, we used *CellChat* (Majidian et al. 2025) to infer intercellular signaling probability based on the expression of secreted ligands and their cognate receptors across cell types (Figure 2B). By analyzing the signaling inputs received by rock dove gPGCs from surrounding cell populations, we aimed to identify candidate pathways and factors that could inform the optimization of *in vitro* culture conditions. This analysis revealed that growth-related communication was dominated by midkine (MDK) and pleiotrophin (PTN) signaling through receptors such as nucleolin (NCL), LRP1, and syndecans (SDC1/4), together accounting for the majority of predicted growth-supporting input. Additional contributions came from canonical WNT ligands (WNT4, WNT16) engaging FZD–LRP co-receptors, as well as EGF–EGFR, and KITL–KIT. Retinoic acid signaling components were also detected, suggesting a balance between pro-proliferative and differentiation cues within the gonadal niche (Figure 2C).

Importantly, rock dove gPGCs at stage HH28 did not express the differentiation marker *FOXL2L* (Figure 1B), suggesting that the gonadal PGCs maintain early germ cell identity and could be further propagated to generate cell lines (Biegler et al. 2025; Liu et al. 2022).

### Osmolality and Growth Factor Requirements for Rock Dove PGCs

We initially employed the low osmolarity FAOT medium originally formulated for chicken PGCs as the basis for our experiments (Whyte et al. 2015). Whyte and colleagues reported that the osmolality of embryonic chicken blood, through which PGCs migrate, is 260 mOsm/kg—lower than that of most basal media—and therefore adjusted their formulation accordingly. To determine whether a similar adjustment was needed for rock dove, we measured the osmolality of embryonic serum and found it to be highly comparable at 256.7 ± 4.4 SEM mOsm/kg. On this basis, no further modification of the medium’s osmolality was required. However, in FAOT medium, rock dove PGCs differentiated and survived only a few days.

Studies in goose showed that activin A, a component of FAOT, inhibits goose PGC growth whereas BMP4 promotes PGC proliferation (Doddamani et al. 2025). To test whether BMP signaling is more effective than activin A in maintaining rock dove PGC survival and proliferation, cells were cultured in media supplemented with either BMP4 or activin A. Both activin A and BMP4-treated PGC cultures maintained PGC identity after 2 weeks of culture, as determined by RT-qPCR of *DAZL, DDX4*, and *PRDM14* (Figure 3A), however, BMP4-treated cultures exhibited significantly greater cellular outgrowth and proliferation compared to activin A-treated cells (Figure 3B). These findings indicate that BMP signaling may be the preferred pathway for supporting rock dove PGC maintenance. Furthermore, when treating cells with BMP4 while inhibiting the activin A pathway with a selective inhibitor of the TGF**β** type 1 receptors, SB431542, we observed greater numbers of PGCs, identified using GFP-tagged *Lycopersicon esculentin* lectin, which has been shown to bind specifically to PGCs (Iikawa et al. 2024; Supp. Fig. 4).

**Figure 3:**
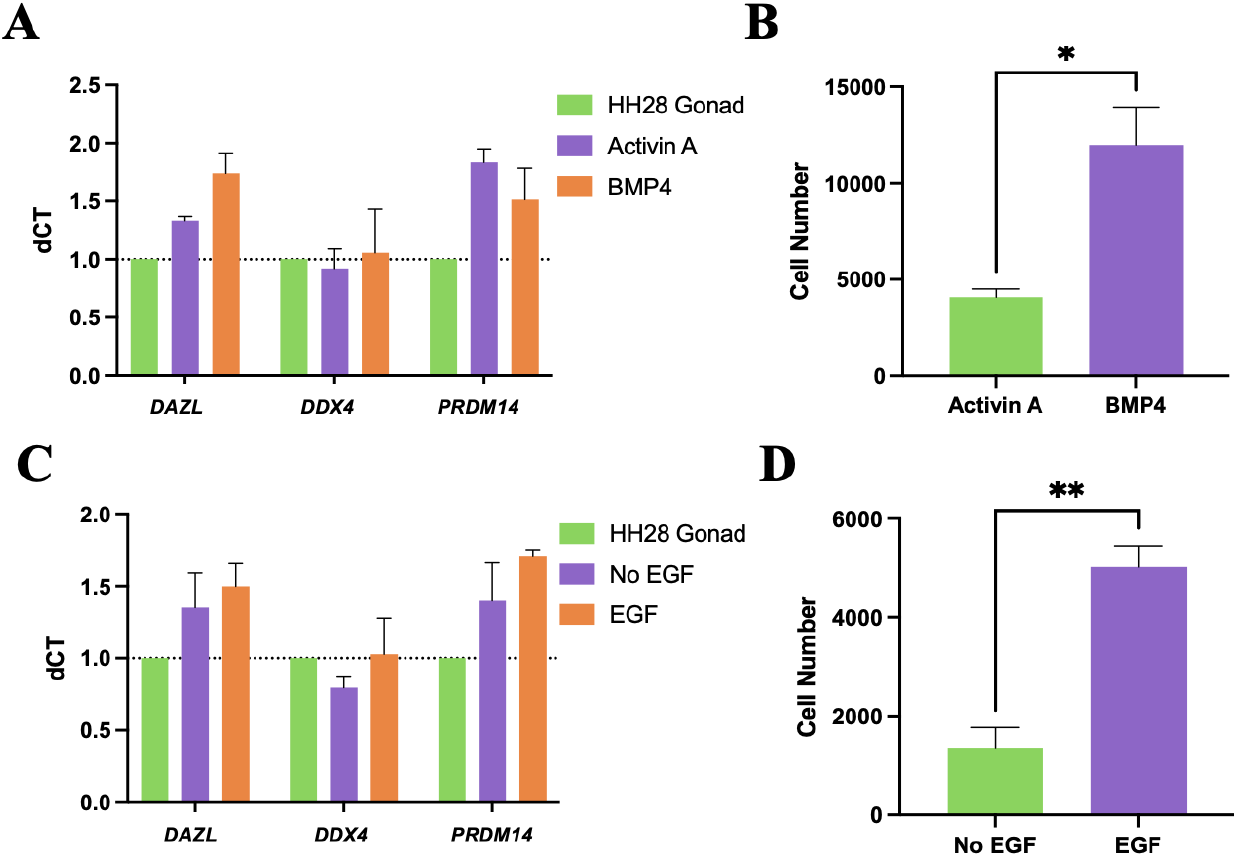
Rock dove PGC proliferation is dependent on BMP4 and EGF signalling. Rock dove gonads were cultured in either activin A or BMP4 containing media. (A) RT-qPCR of PGCs cultured in either activin A or BMP4 containing media. (B) Quantification of PGCs in culture. (C) RT-qPCR of PGCs cultured for 2 weeks with or without 10 ng/ml EGF. A sample of cells were imaged and quantified (D). Error bars, SEM. * p<0.05, **p<0.01.

Our scRNA-seq data showed that EGFR is expressed in rock dove PGCs (Figure 2A). To assess the importance of EGF signaling, rock dove PGCs were cultured in media containing EGF. These cultures maintained PGC identity, shown through *DAZL, DDX4*, and *PRDM14* expression after 14 days in culture (Figure 3C). Furthermore, the addition of EGF resulted in increased cell number compared to cultures lacking EGF (Figure 3D), indicating that EGF signaling is beneficial for PGC proliferation.

### Insulin/IGF Signaling is Essential for PGC Propagation

B27 supplement, a key component of FAOT and many avian PGC culture media formulations, typically contains insulin at micromolar concentrations—significantly higher than the physiological levels found *in ovo*, which are in the picomolar range (Lu et al. 2007). Elevated insulin levels have been shown to negatively impact insulin and IGF receptor function, ultimately inhibiting downstream insulin signaling (Catalano et al. 2014). To examine the effects of insulin on PGC outgrowth, we tested a modified version of B27 supplement lacking insulin, and assessed expression of canonical germline markers *DAZL, DDX4*, and *PRDM14*. While insulin-free conditions preserved PGC identity, they did not result in a significant increase in cell number (Figure 4A–B), and cultures could not be maintained beyond two weeks (data not shown), suggesting that some insulin signaling is necessary for long-term proliferation.

**Figure 4:**
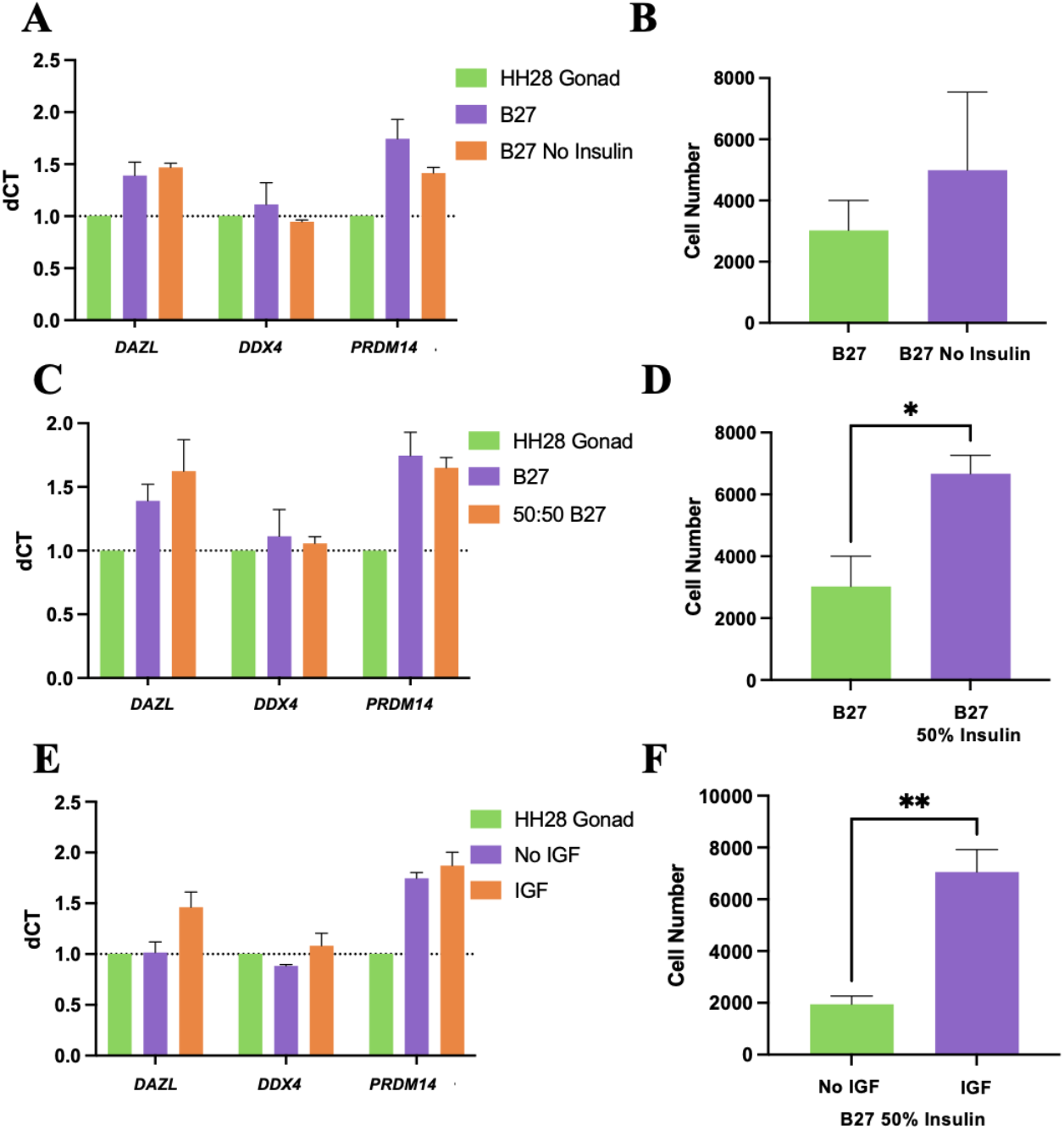
Insulin and IGF signaling are important for rock dove PGC survival and proliferation. RT-qPCR of PGC markers and cell number quantification of cells cultured for two weeks in media containing (A-B) B27 with or without insulin, (C-D) B27 with insulin or a 50:50 mixture of B27 with and without insulin (B27 50% Insulin), and (E-F) cultured with B27 50% Insulin with or without 10 ng/ml IGF. Error bars, SEM. * p<0.05, **p<0.01.

To address this, we reduced the insulin concentration by mixing standard B27 with insulin-free B27 in a 1:1 ratio (B27 50% Insulin). This modification supported increased cell proliferation compared to B27 supplement while maintaining germline identity (Figure 4C–D). Furthermore, addition of IGF under reduced-insulin conditions (B27 50% Insulin) enhanced PGC expansion when compared to IGF-free conditions, without compromising the expression of key germline markers (Figure 4E–F).

### Retinoic Acid Receptor Blockade Enables PGC Expansion

Retinoic acid signaling is known to drive PGC differentiation (Bowles et al. 2006; Anderson et al. 2008). Vitamin A, a precursor of retinoic acid, is another standard component of B27 supplement. To reduce differentiation of PGCs in culture, we tested B27 specially formulated without vitamin A. Cells cultured without vitamin A maintained PGC markers and a comparable growth rate at 14 days in culture compared to control media (Figure 5A-B), however, these cells did not continue to proliferate beyond 2-3 weeks leading to differentiation and cell death. Conversely, inhibiting the retinoic acid receptor with BMS493 improved PGC outgrowth and cell survival while maintaining germline identity when using B27 with vitamin A (Figure 5C-D).

**Figure 5:**
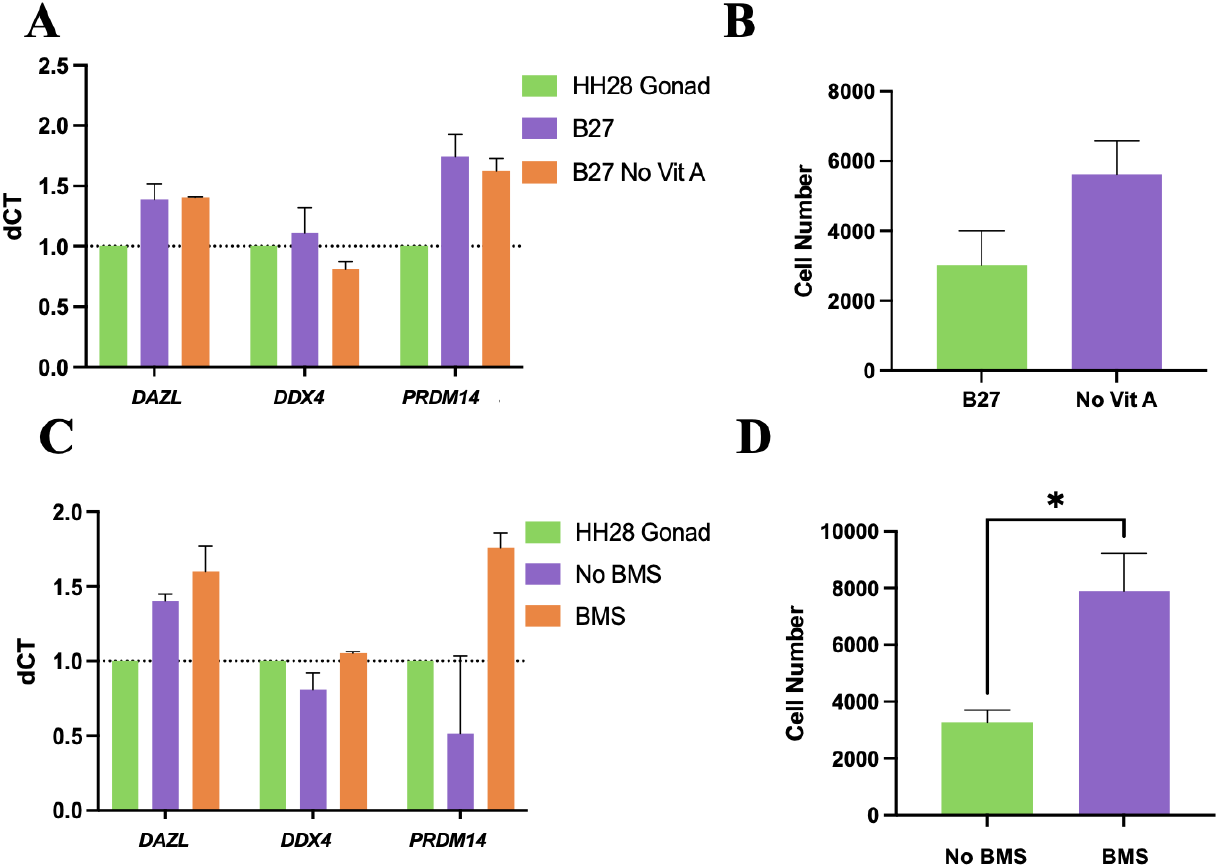
Long-term culture of rock dove PGCs requires vitamin A and is improved by retinoic acid receptor blockade. RT-qPCR of PGC markers and cell number quantification of cells cultured for two weeks in media containing (A-B) B27 with or without vitamin A and (C-D) with or without 5 µM BMS493. Error bars, SEM. * p<0.05.

### Pleiotrophin promotes PGC proliferation

CellChat analysis suggested that the heparin-binding proteins midkine and pleiotrophin provide important intercellular signals to PGCs. Midkine has previously been shown to promote proliferation in several cell types, including PGCs (Shen et al. 2012). In our experiments, both factors supported maintenance of germline identity (Figure 6A, C), but only pleiotrophin at 10 ng/ml significantly increased cell number. In contrast, high concentrations (100 ng/ml) of either midkine or pleiotrophin reduced cell numbers compared to control and low-dose treatments (Figure 6B, D).

**Figure 6:**
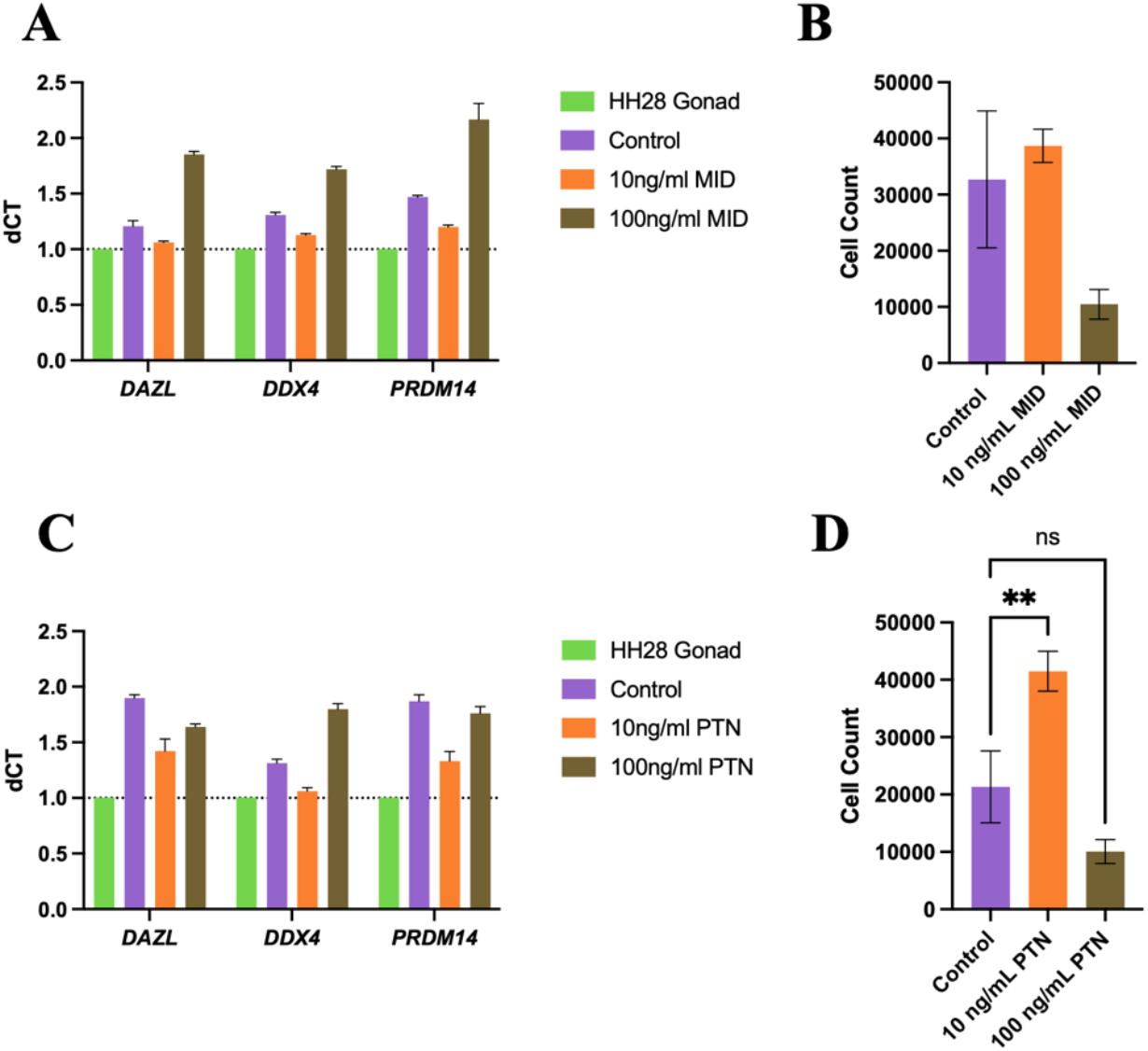
Pleiotrophin but not midkine increases rock dove primordial germ cell proliferation. RT-qPCR of PGC markers and cell number quantification of cells cultured for two weeks in media containing (A-B) midkine or (C-D) pleiotrophin. Error bars, SEM. ** p<0.01.

### Cultured Rock Dove PGCs Express PGC Markers and Migrate to the Embryonic Gonad

After establishing culture conditions that support PGC culture (Figure 7A), we additionally assessed expression of markers such as *SOX2*, a key regulator of pluripotency (Avilion et al. 2003), and *CXCR4*, a receptor required for migration to the gonad (Doitsidou et al. 2002), and confirmed the absence of *FOXL2L*, a marker of germ cell differentiation (Supp. Fig. 5). We also determined that these cells have a calculated doubling rate of ∼35 hours (Figure 7B), similar to that reported for chicken PGCs (Kinoshita et al. 2024; Altgilbers et al. 2021).

**Figure 7:**
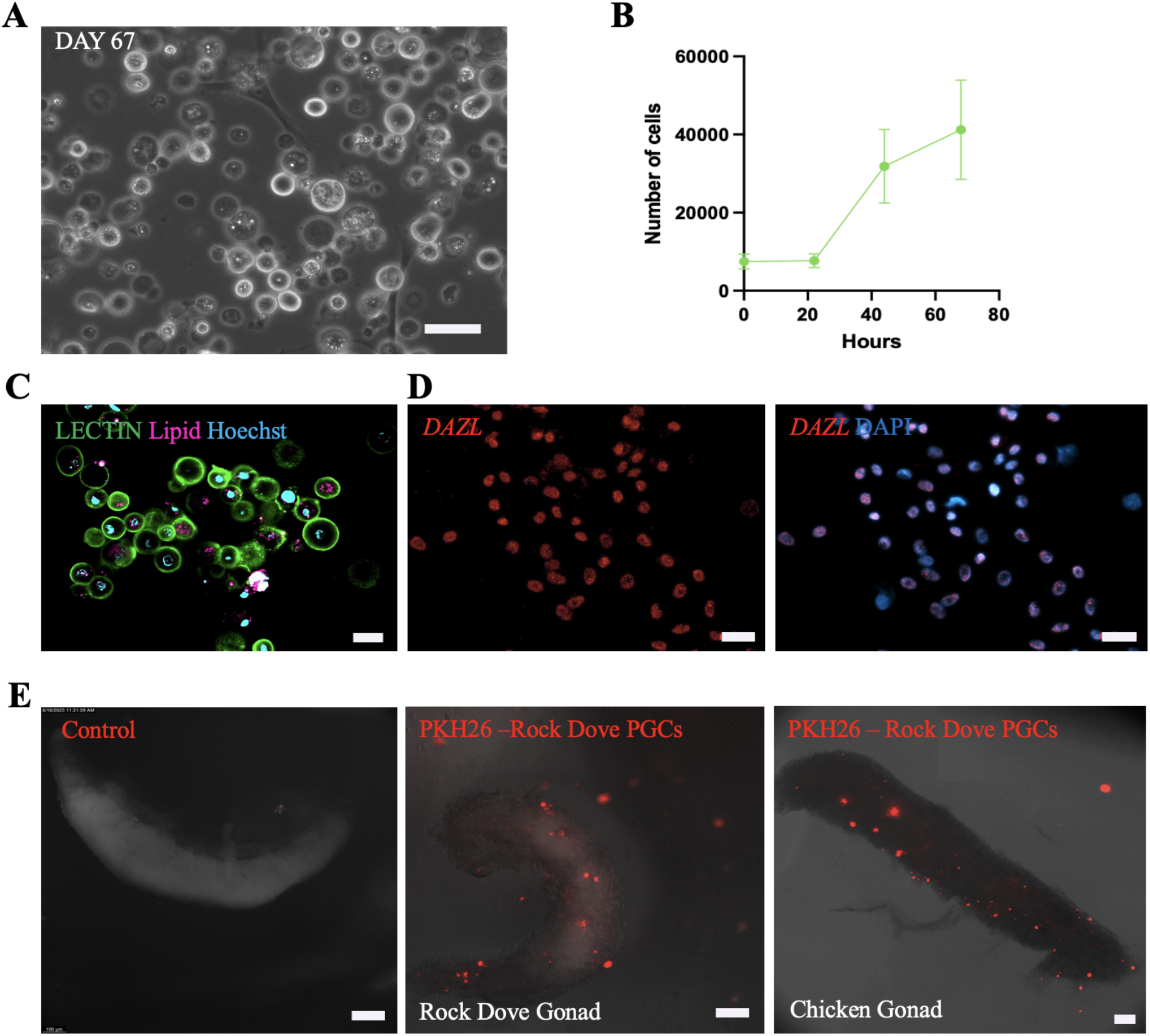
Cultured rock dove PGCs migrate to the embryonic gonad. (A) Brightfield image of rock dove PGCs cultured for 67 days. (B) Cell counts every 24 hours quantified using CellaVista. Calculated average doubling rate of 35.4 hrs. Error bars = SEM. (C) Live cell imaging of cultured rock dove PGCs stained with tomato lectin (green), LipidSpot (red), and Hoechst (blue). (D) Immunocytochemistry of cultured rock dove PGCs stained with DAZL (red) and DAPI (blue). (E) Stage HH28 rock dove and chicken embryonic gonad with migrated cultured rock dove PGCs stained with PKH26. Scale bar in A= 50 µm, B-C = 20 µm, Scale bar in C = 100 µm.

To further evaluate culture purity, we performed live cell imaging of PGCs labeled with GFP-tagged *Lycopersicon esculentin* lectin and a lipid dye to visualize lipid granules, a characteristic of PGCs. We found a highly pure population of lectin-positive cells with accumulated lipid granules and few non-stained cells (Figure 7C). Immunocytochemistry using an anti-DAZL antibody showed that the majority of DAPI-positive nuclei also expressed DAZL, supporting the conclusion that our cultures contained a highly pure PGC population (Figure 7D).

We next evaluated the functional capacity of cultured rock dove PGCs to migrate to the embryonic gonad. PKH26-fluorescent stained rock dove PGCs were injected into rock dove HH15 embryos and embryos were incubated for 4 days. PKH26-positive cells were detected within the gonadal tissue of injected embryos at stage HH28. Additionally, cultured rock dove PGCs showed the ability to migrate to the gonads of chicken embryos (Figure 7E), indicating cross-species migratory competency.

## Discussion

Previous studies have described variations in culture media that support the long-term *in vitro* growth of chicken and goose PGCs (van de Lavoir et al. 2006; Whyte et al. 2015; Doddamani et al. 2025). In our initial experiments, both chicken FAOT and goose PGC media failed to sustain long-term proliferation of rock dove PGCs, highlighting the need for species-specific optimization. Here, we demonstrate an omics-guided approach to developing culture conditions tailored to rock dove PGCs by integrating scRNA-seq profiling of embryonic gonadal cells. This strategy enabled us to identify key growth factor and signaling requirements, resulting in a more effective and species-appropriate medium than protocols adapted from other avian species (Figure 8).

**Figure 8:**
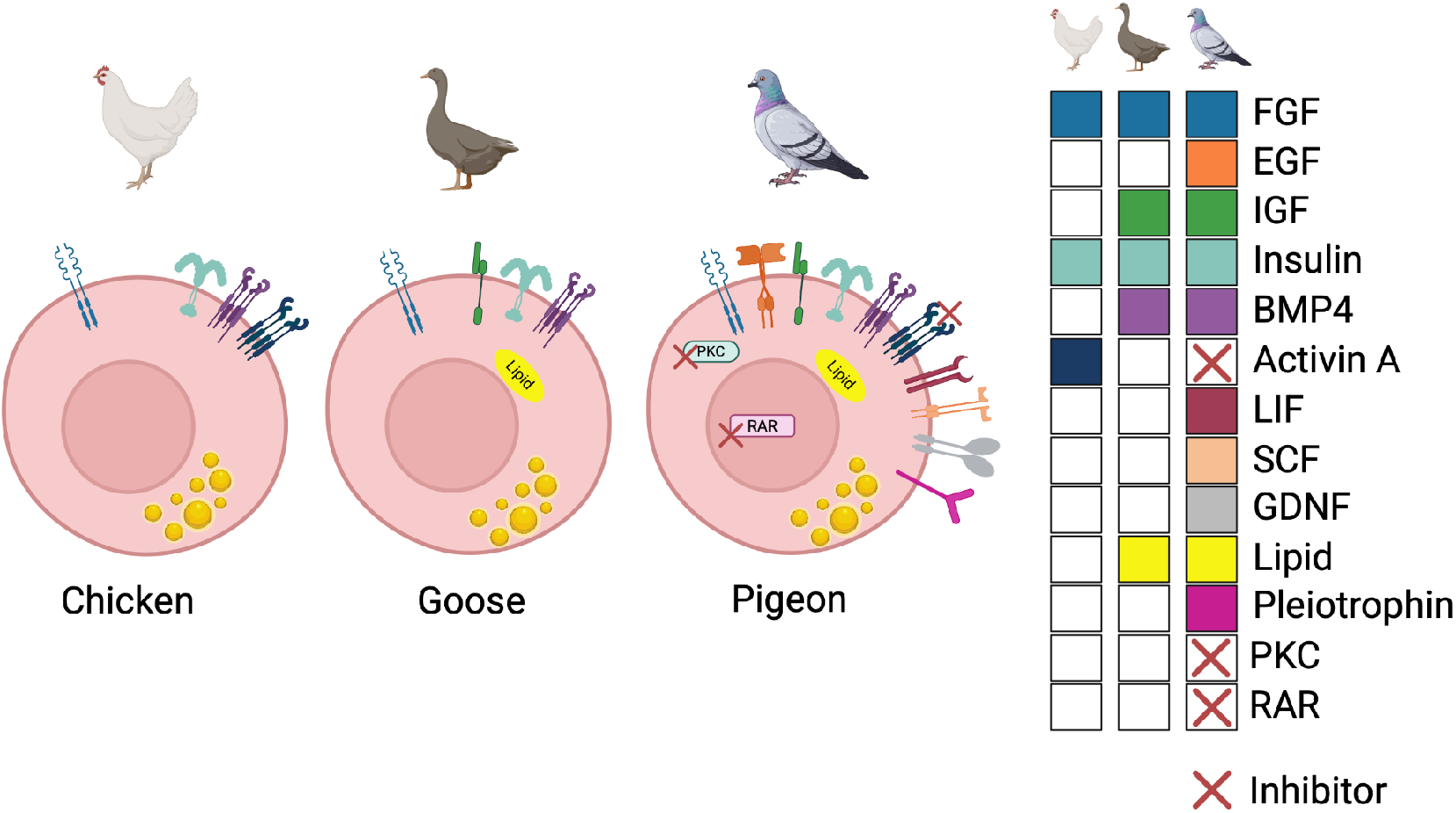
Schematic of differences between chicken, goose, and rock dove PGC growth factor requirements. List of growth factors and their respective receptors or pathways in either chicken (left), goose (middle), or rock dove (right) that are required for PGC survival and proliferation. Red ‘X’ signifies inhibition of the pathway/receptor. Created in https://BioRender.com

Our scRNA-seq data revealed high expression of *ACVR2A/B*, suggesting potential responsiveness to Activin/TGF-β signaling. Consistent with findings in goose PGCs, activin A inhibited rock dove PGC proliferation—despite its supportive role in chicken PGC culture (Whyte et al. 2015). This contrast reinforces the emerging view that avian PGCs exhibit species-specific sensitivities to signaling environments. Nevertheless, similarities are beginning to emerge between avian species such as the requirement for FGF, insulin/IGF, and BMP signaling. Future studies will be required to test if the rock dove media formulation would support PGC cultures of other avian species or enhance established media formulations for chicken and goose.

The high expression of retinoic acid receptors alongside low *FOXL2L* expression suggested that rock dove PGCs remained undifferentiated but might be susceptible to premature differentiation if exposed to retinoic acid signaling. Removing vitamin A from the medium impaired cell survival, but pharmacologic inhibition of retinoic acid receptors (with BMS493) in the presence of vitamin A supported long-term proliferation. This suggests that vitamin A may serve alternative, RAR-independent roles in PGC physiology. Further research is needed to pinpoint these alternative pathways in PGCs.

We also observed high expression of insulin/IGF pathway receptors but found that the supraphysiologic insulin concentrations in standard B27 supplement negatively affected proliferation—possibly due to receptor downregulation, as shown in previous studies (Biadgo et al. 2020). Reducing insulin concentrations while supplementing with IGF supported cell proliferation and preserved germline identity, illustrating the importance of fine-tuning media composition, even within broadly conserved pathways.

CellChat profiling showed that two heparin-binding proteins, midkine and pleiotrophin, may provide important signaling to gonadal PGCs. Midkine has previously been shown to increase PGC proliferation through inhibiting DAZL in non-avian models (Shen et al. 2012). Interestingly, we found that these two similar proteins did not alter DAZL expression and only pleiotrophin increased PGC numbers. Although midkine and pleiotrophin share 50% homology (Weng and Liu. 2010) and share many downstream signaling pathways such as PI3K/AKT and MAPK/ERK signaling, pleiotrophin emphasizes Wnt/β-catenin stabilization via PTPRζ inhibition (Aller et al. 2025; Muramatsu. 2002), suggesting this pathway specifically may be important for PGC proliferation.

Together, these optimizations enabled us to maintain rock dove PGCs in culture for over 60 days. Importantly, cultured cells retained functional competency, as demonstrated by their ability to migrate to the gonads of both pigeon and chicken embryos, confirming their identity and developmental potential. Migration efficiency may be further enhanced through the use of germline-ablated surrogate hosts, which reduce competition from endogenous PGCs (Ballantyne et al. 2021; Taylor et al. 2017; Woodcock et al. 2019). Future work will focus on assessing germline transmission capacity, which remains the definitive test of functional competence and will be essential for downstream applications.

Establishing reliable culture systems for PGCs from Columbidae is especially timely. This family, which includes over 300 species, is among the most threatened bird lineages globally (Walker 2007). The IUCN Red List currently identifies 68 pigeon and dove species as globally threatened, 14 of which are critically endangered or extinct in the wild (IUCN 2025). Our work lays a foundation for germline preservation, assisted reproduction, and genetic rescue in this family (van Oosterhout et al. 2025). More broadly, it demonstrates a scalable, data-driven strategy for extending germline technologies beyond chickens to other avian taxa—an essential step for expanding the application of artificial reproductive technologies to conservation.

## Materials and Methods

### Animal Welfare

All procedures involving avian embryos were conducted in compliance with institutional and federal guidelines on avian embryo use. According to U.S. regulatory and NIH Office of Laboratory Animal Welfare (OLAW) interpretations, embryos of egg-laying species are not considered live vertebrate animals prior to hatching. All chicken and rock dove embryos were incubated no longer than 7 days.

### PGC Collection

Fertile rock dove eggs were obtained from local, private breeders and incubated for 6 days to the equivalent of stage HH28. Gonads were dissected out, cut in two, and each pair were plated in TC-treated 24 well plates with 600 µL of calcium-free medium per well for the first 24 hours to facilitate PGC migration out of the gonadal tissue. The next day, the media was changed to include 0.15 mM CaCl_2_ (Sigma: C34006). One third of the media was replaced every 48 hours thereafter. Cultures were grown for a minimum of two weeks and assayed for proliferation and cell number.

### Rock Dove Primordial Germ Cell Culture

Cultures were maintained in custom Knockout-DMEM base medium (Thermo Fisher; Whyte et al. 2015) supplemented with Glutamax (Gibco; 17504044), NEAA (Gibco; 11140035), Sodium Pyruvate (Gibco; 11360039), 0.5X B27 minus insulin (Thermo Fisher; A1895601), 0.5X B27-no vit A (Thermo Fisher; 12587010), 0.4% chicken serum (Sigma; C5405), 50 µg/ml ovotransferrin (Sigma; C7786), 0.2% ovalbumin (Sigma; A5503), 0.1 mg/ml Na Heparin (Sigma; H3149), 4 ng/mL FGF1 (PeproTech; 100-17A), 4 ng/mL FGF2 (PeproTech; PHG0369), 5 ng/mL BMP4 (R&D Systems; 314-BPE), 10 ng/mL IGF1 (PeproTech; 100-11), 10 ng/mL EGF (PeproTech; AF-100-15), 1 µg/mL cholesterol (Sigma; C8667), 50 µM β-mercaptoethanol (Thermo Fisher), 0.15 mM CaCl_2_, 50 ng/mL cLIF (Kingfisher; RP1395C), 0.25 µM blebbistatin (Sigma; B0560), to 350 mg/dL glucose (Thermo Fisher; A2494001), 10 µM SB431542 (Tocris; 1614), 1:1000 lipid mix (Sigma; L0288), 5 µM BMS493 (Tocris), 25 ng/mL cSCF (Kingfisher; RP1555C), 5 ng/mL GDNF (PeproTech), and Pen/Strep (Gibco; 15070-063). Custom Knockout-DMEM was prepared to an osmolality of 250 mOsm/kg, with 12.0 mM glucose and without calcium. Cultures were maintained at 37°C in a humidified incubator with 5% CO_2_.

### scRNA-seq

#### PGC Collections for scRNA-seq from Gonads

Fertile rock dove eggs were incubated for 6 days to the stage equivalent of HH28. Gonads were dissected out and pooled in room temperature TrypLE™ Express Enzyme (Gibco, 12604201). Collections were incubated at 37°C for 20 minutes with manual dissociation by gentle pipetting in 5-minute intervals. Dissociated gonadal cells were pelleted, resuspended in PGC media and counted using the CellDrop FL automated cell counter with AO/PI staining (DeNovix). Cells were diluted in GMP grade Stem Cell Banker (TAKARA Bio, 11922) and frozen for later processing. Gonads from 6 rock dove embryos were combined, yielding ∼1.65×10^5^ total cells with a 94.1% viability.

#### Sample preparation and scRNA-seq library

Cryopreserved primordial germ cells of two *Columba livia* samples (PGC; n = 2) were shipped in cryopreservation media to the University of Connecticut. All samples were processed using the Fluent Biosciences PIPseq V T20 3’ Single Cell RNA kit (revision 1.5), following the manufacturer’s protocol for cryopreserved cells with modifications as detailed below. Cryovials were thawed in a 37°C water bath until approximately 70% thawed. The cell suspension was then transferred using a wide-bore P1000 pipette tip into a 15 mL conical tube containing pre-warmed DMEM. Samples were gently mixed by inversion (5x) and centrifuged at 200 × *g* for 5 minutes at room temperature using a swinging-bucket centrifuge. Supernatant was removed, and the cell pellet was resuspended in 500 µL of pre-warmed Cell Suspension Buffer.

Cell viability and count were assessed using the ThermoFisher Countess Cell Counter with Trypan Blue staining. Total viable cell counts ranged from 45,000 to 500,000 cells per sample. Cell suspensions were normalized to 5,000 cells/µL, and 8 µL (equivalent to 40,000 cells) were loaded into each PIPseq tube. Following cell encapsulation, the Cell Lysis Program was performed on a dry bath. After lysis, samples were held at 20°C overnight. During cDNA amplification for quality control, 11 PCR cycles were used to accommodate the low input of starting material. cDNA quantity was assessed using the dsDNA High Sensitivity Assay for Qubit 3.0 (Life Technologies, Carlsbad, CA, USA). Fragment size distribution was evaluated using the Agilent Tapestation 4200 D5000 High Sensitivity assay (Agilent Technologies, Santa Clara, CA, USA).

Illumina library preparation was subsequently performed. Fragmentation was conducted for 6 minutes based on average cDNA fragment sizes. Dual index mixes (Fluent Biosciences) were used for the Sample Index PCR, with 13 cycles of amplification applied due to the low input material. Final libraries were validated for length and adapter dimer removal using the Agilent TapeStation 4200 D1000 High Sensitivity assay (Agilent Technologies, Santa Clara, CA, USA) then quantified and normalized using the dsDNA High Sensitivity Assay for Qubit 3.0 (Life Technologies, Carlsbad, CA, USA).

Sample libraries were prepared for Illumina sequencing by denaturing and diluting the libraries per manufacturer’s protocol (Illumina, San Diego, CA, USA). All samples were pooled into one sequencing pool, equally normalized, and run as one sample pool across the Illumina NovaSeq 6000 using version 1.5 chemistry. Final loading concentration was 0.6nM and 1% PhiX used. Target read depth of 20,000 cDNA reads was achieved per cell, per sample with paired end reads. Each T20 library targeted 800M total PE reads. Read 1, Index 1 and 2, and Read 2 sequencing cycles were as follows: 45, 10, 10, 72.

#### scRNA-seq bioinformatics

Raw count matrices for rock dove gonadal cells were generated using the PIPseeker pipeline v3.3.0 (Fluent Biosciences). In brief, sequenced reads were mapped to genes by matching their genomic coordinates to the *Columba livia* genome reference (NCBI reference assembly bColLiv1.pat.W.v2; accession GCF_036013475.1). Reads with the same molecular identifier (MI) were considered duplicates and collapsed into a single count. The resulting data were stored in a sparse matrix, with each entry representing the count for a specific cell barcode–gene pair. Barcodes were classified as either cell-associated PIPs or background PIPs based on a barcode rank (“knee”) plot, which ranks barcodes by their total transcript counts. The final output consisted of a filtered count matrix (matrix.mtx.gz, barcodes.tsv.gz, and features.tsv.gz) with a moderate sensitivity of 3. Data were imported into R (v4.3.2) and processed using the Seurat package (v5.0.3; Hao et al. 2023). Gene annotations were curated to replace *LOC* pigeon gene identifiers with corresponding *Gallus gallus* orthologs gene symbols (NCBI reference assembly bGalGal1.mat.broiler.GRCg7b; accession GCF_016699485.2) based on an orthology assignment by fastOMA (Butler et al. 2018).

For each cell, quality control (QC) metrics were computed, including the number of detected genes (nGene), total unique molecular identifiers (nUMI), the ratio of genes to UMIs (log_10_-transformed), and the proportion of mitochondrial transcripts (mitoRatio). Cells were retained if they contained >1,000 UMIs, >500 detected genes, and <20% mitochondrial RNA content, thresholds chosen based on inspection of QC metric distributions. Following filtering, data were normalized using SCTransform (v2; Hafemeister and Satija, 2019), regressing out variation associated with cell cycle phase, mitochondrial proportion, and sequencing depth.

Highly variable genes were identified using variance-stabilizing transformation, and principal component analysis (PCA) was performed on the top 2,000 features. The first 25 principal components were used to compute a Uniform Manifold Approximation and Projection (UMAP) embedding (Becht et al. 2019). Shared nearest neighbor (SNN) graph-based clustering was performed (*FindNeighbors* and *FindClusters*), with a primary resolution of 0.4 selected for downstream analyses after resolution exploration. Cell type annotations were assigned by examining the top 50 differentially expressed genes in each cluster and identifying canonical marker genes supported by literature and previous avian PGC studies (Biegler et al. 2025). These annotations were then cross-referenced with published avian single-cell datasets, and clusters corresponding to contaminant renal cells or low-identity cycling populations were removed.

#### CellChat

To reconstruct ligand–receptor–mediated intercellular signaling networks from our gonadal single-cell transcriptomes, we employed the R package CellChat (v2; Majidian et al. 2025), which integrates curated interaction databases with cell-type–specific expression profiles. We first loaded Seurat objects containing harmonized scRNA-seq data for chicken and rock dove gonads and converted them into CellChat objects. We assigned the human ligand–receptor database (CellChatDB.human) to each object and filtered the expression data to retain only signaling genes. We then identified over-expressed ligands, receptors, and their interactions using the *identifyOverExpressedGenes* and *identifyOverExpressedInteractions* functions, respectively. Next, we computed communication probabilities using a trimmed-mean model and filtered for interactions supported by at least ten cells. Pathway-level interaction probabilities were then derived via *computeCommunProbPathway*, and species-specific interaction tables were extracted from aggregated interaction networks. Finally, we ranked these interactions by interaction probability and filtered only those involving primordial germ cells.

### RT-qPCR

Total RNA was extracted using the RNeasy Mini Kit (Qiagen; 74104) and RNA yield was quantified using a NanoDrop spectrophotometer. cDNA was synthesized using the SuperScript IV First Strand Synthesis kit (Invitrogen; 18091050). qPCR reactions were prepared with PowerUp SYBR Green Master Mix (Applied Biosystems; A25742). Each transcript of interest was amplified with primers (Sup. Table 1) designed to span exon-exon junctions and target all known transcript variants for each gene. qPCR reactions were run using a QuantStudio 7 system (Applied Biosystems) with the following settings: 50°C (2 min), 95 °C (2 min), 95 °C (15 s), 55 °C (15 s), 72 °C (1 min) (return to step 3 for 40 cycles). A continuous melt curve analysis was performed on each sample using the following settings: 95°C (15 s), 60 °C (1 min), 95 °C (15 s). The *GAPDH* housekeeping gene was used as an internal control to normalize gene expression data. Results were analyzed using the Applied Biosystems Relative Quantification analysis online application (version 2021.1.1-Q1-21-build11) to determine delta CT (dCT).

### Immunocytochemistry (ICC) Staining

Cultured PGCs were spun at 500 rpm for 5 minutes onto charged microscopy slides (Fisher; 1255015) using a Cytospin 4 centrifuge (Thermo Scientific). Cells were fixed with 4% paraformaldehyde and permeabilized using 0.1% Triton X-100 for 10 minutes. Blocking was performed with 5% goat serum for 30 minutes at room temperature. Slides were incubated overnight at 4°C in a humidified chamber with primary anti-DAZL antibody (Abcam EPR21028) at a 1:1000 dilution. A donkey anti-rabbit Alexa Fluor 488 secondary antibody was used at 1:1000 dilution for 1 hour at room temperature in the dark. Slides were mounted using Prolong Gold Antifade Mount with DAPI (Thermo Fisher; P36931) and imaged using a Nikon AX-NSPARC resonant scanning confocal equipped with a sCMOS camera using a 20X (Plan Apo, NA 0.45) objective.

### Cell Quantification

Cells were plated in a CytoOne 96 well plate (USA Scientific; CC7682-7596) and imaged on a CellaVista Imaging System (Synentec). Cells were analyzed using brightfield images based on size, roundness, and contrast using the CellaVista imaging software.

### Live Cell Imaging

Cells were collected in a 1.5 mL tube, washed with DPBS and centrifuged at 300xg for 5 minutes. Cells were incubated with *Lycopersicon esculentum* Lectin DyLight 488 (Thermo Fisher; L32470) at 1:1000 dilution, LipidSpot 610 (Biotium; 70069) 1:1000 dilution, and 10 µM Hoechst 33342 (Thermo Fisher; 62249) diluted in DPBS for 30 minutes at 37°C in a humidified incubator. Cells were washed 3 times with DPBS, plated on an Ibidi 12 chamber slide (Ibidi; 81201) and imaged using a Nikon AX-NSPARC resonant scanning confocal equipped with a sCMOS camera and a 20X objective (Plan Apo, NA 0.45).

### Migration Assays

Cultured PGCs grown for a minimum of 30 days were labeled with PKH26 dye (Sigma; PKH26GL) and resuspended at 5,000 cells/µl in custom Knockout-DMEM with 0.075uM CaCl_2_ and 0.01% sterile-filtered fast green (Sigma; F7252-5G) in PBS (for injection visualization). Approximately 10,000 PGCs (∼2µl) were injected into the dorsal aorta of chicken or rock dove embryos at stage (or stage equivalent of) HH13-

16. Eggs of injected embryos were sealed with tape and incubated for an additional 4 days to allow PGCs to migrate to the gonad. Gonads were dissected at stage HH28 and imaged using a Leica M205FCA fluorescent microscope equipped with a Leica K5 camera. PGC migration was confirmed by fluorescent visualization in the dissected gonads and compared to un-injected, stage matched controls.

## Data Availability

The raw sequencing data can be accessed through the NCBI Sequence Read Archive under BioProject accession number PRJNA1321384, SRA Study SRP618147, and SRA Run accession number SRX30391834, which have been made available to reviewers.

## Acknowledgements

The authors thank Matt Biegler for discussions on PGC culture and analysis. We thank the Center for Genome Innovation (in the Institute for Systems Genomics) and Bo Reese for sequencing support.

## Author Contributions

MWN: Conceptualization; Formal analysis; Investigation; Methodology; Project Administration; Supervision; Validation; Visualization; Writing-original draft.

LM: Data curation; Formal analysis; Methodology; Software; Writing - original draft.

EAB: Investigation; Methodology; Resources; Validation; Visualisation; Writing-original draft. EM: Investigation; Methodology; Resources; Writing-original draft.

AS: Resources; Writing - review & editing. MM: Resources.

WNF: Investigation; Validation; Writing - original draft; Writing - review & editing. SKN: Resources. Writing - review & editing.

KS: Investigation; Resources.

NA: Data curation; Formal analysis; Methodology; Software. SJH: Resources; Methodology.

NMT: Resources; Methodology.

RJO: Supervision; Resources; Methodology; Writing - review & editing. KI: Writing – review & editing

MJM: Writing – review & editing

OF: Project administration; Supervision; Writing - review & editing.

AK: Conceptualization; Project administration; Supervision; Writing - original draft; Writing - review & editing.

