## Supplemental Data for "Long-Term Rock Dove (*Columba livia*) Primordial Germ Cell Culture: A Tool for Avian Conservation"

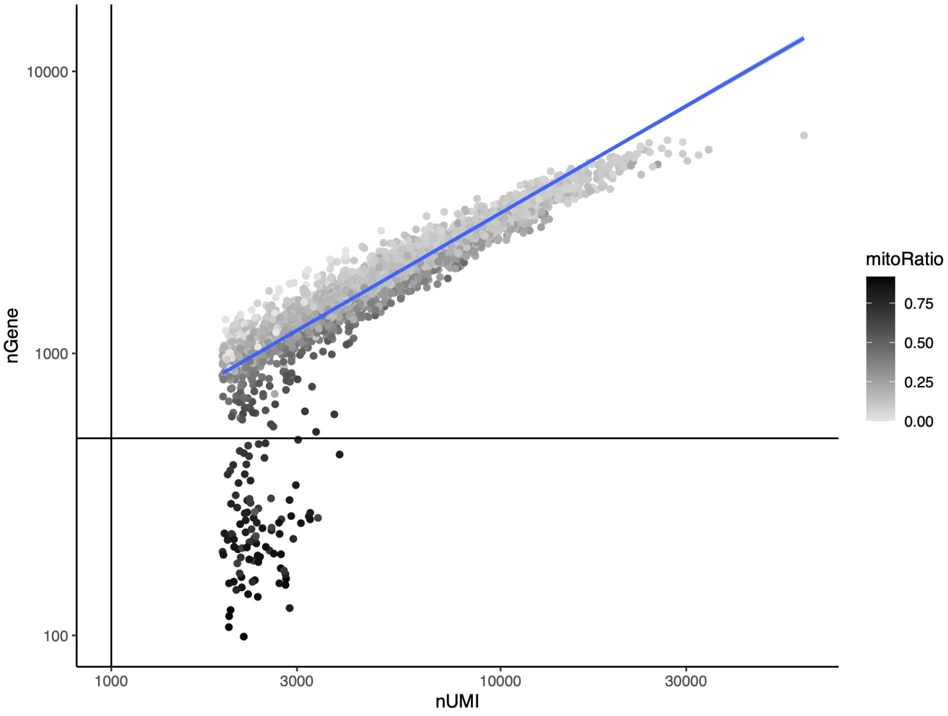


**Sup. Fig. 1: Pre-filtering QC.**
Scatter plot of the number of genes (nGene) vs. library size (nUMI) for all cells. Point shade = mitochondrial fraction; blue line = global trend; horizontal line = nGene cutoff; vertical line = nUMI cutoff. Cells below the cutoff or with high mitochondrial content were excluded.

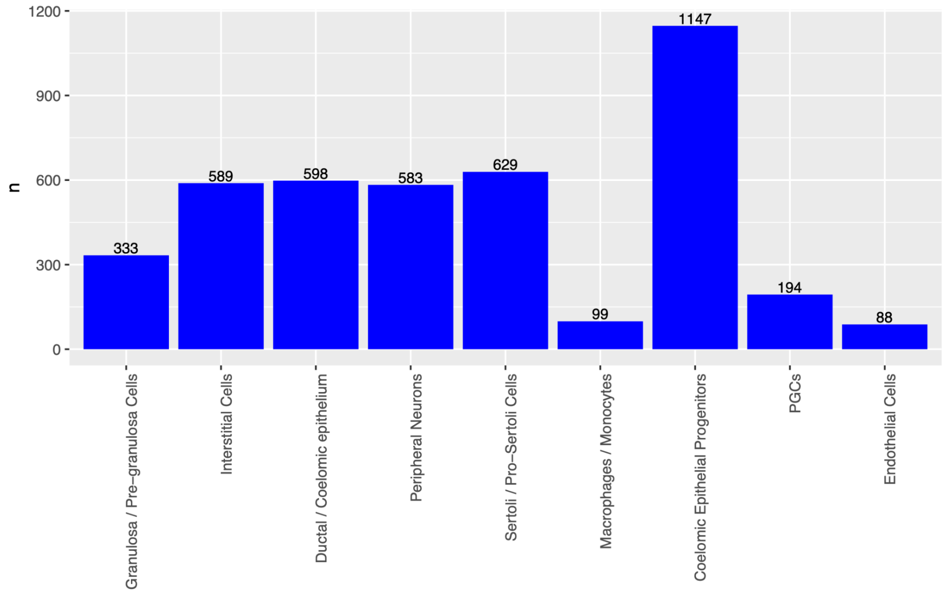


**Sup. Fig. 2: Cell counts.**
Bar chart of number of cells per annotated type; numbers above bars give exact counts.

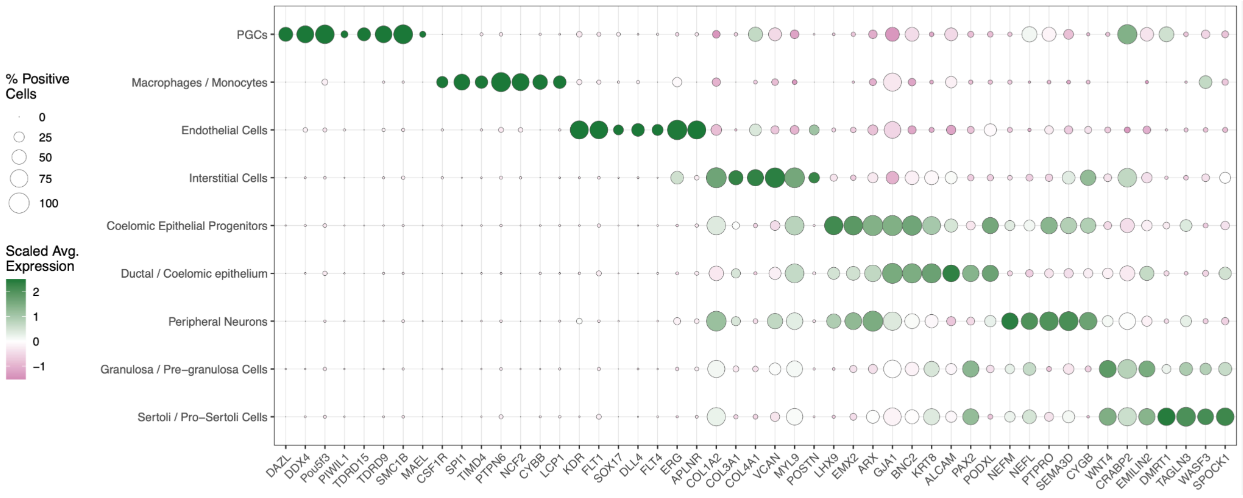


**Sup. Fig. 3: Marker genes by cell type.**
Dot plot of canonical markers across annotated cell types. Dot size = % positive cells; color = scaled mean expression. Used to support cell-type assignments.

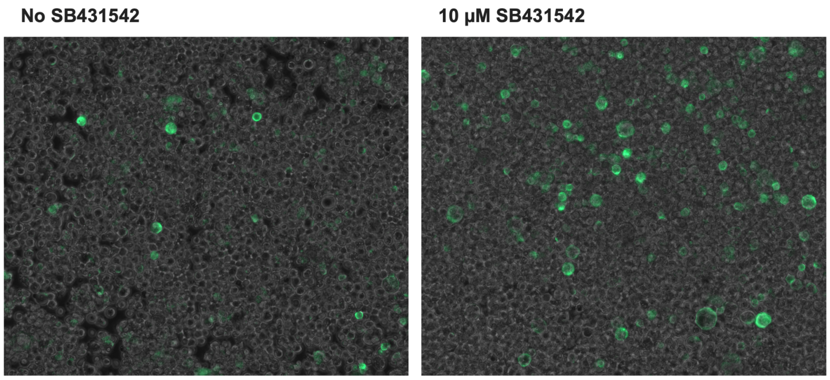


**Sup. Figure 4: Live cell imaging of lectin-stained rock dove gonadal PGC culture.**
Imaging of 2-week PGCs cultured containing BMP4 with or without the TGF**β** inhibitor SB431542.

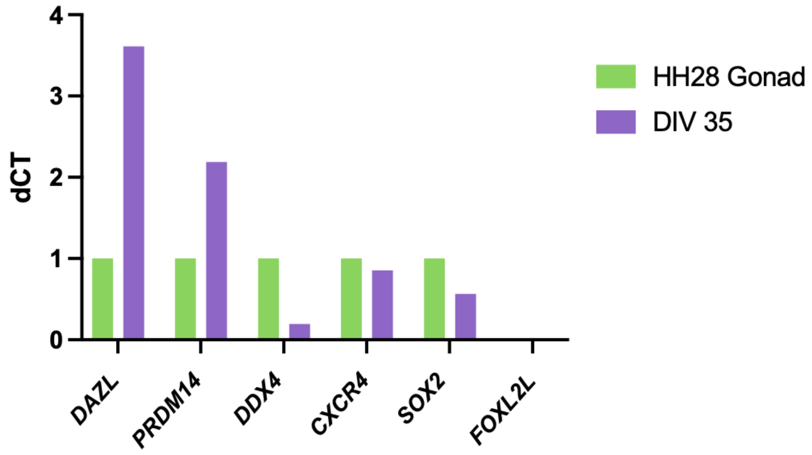


**Sup. Figure 5: RT-qPCR of day 35 cultures**
RT-qPCR panel of canonical PGC markers from day 35 cultures (DIV 35). Cultures express all canonical PGC markers and lack expression of the germ cell differentiation marker *FOXL2L*.

**Sup. Table 1: RT-qPCR Primers**
Primers used for RT-qPCR experiments.

| | **Forward** | **Reverse** |
|---|---|---|
| DAZL | TCCATACAGTTCACCGGCTT | GCCACTGTGGTTGAACCTGAT |
| DDX4 | CTGGAGAAGTTCAGAGGCTGG | GCTGAACATCACTGCAAGCTC |
| PRDM14 | GGCAAAGTGGTCAACACCAG | GCCCGTATTCAAAGATCTCCCA |
| CXCR4 | CAACAACAACTTGAGAGCCCG | TGAACCCAGAGGAGAGATCCA |
| SOX2 | CATTGACGAAGCTAAGCGGC | TCGTCATGGTATTGGTGCCC |
| FOXL2L | TGTCCCTCAACGAGTGCTTC | GACGTTCACCCTGCGGAC |
| GAPDH | GGGTGGTGCTAAGCGTGTTA | GTGCAAGAGGCATTGCTGAC |
